# iTALK: an R Package to Characterize and Illustrate Intercellular Communication

**DOI:** 10.1101/507871

**Authors:** Yuanxin Wang, Ruiping Wang, Shaojun Zhang, Shumei Song, Changying Jiang, Guangchun Han, Michael Wang, Jaffer Ajani, Andy Futreal, Linghua Wang

## Abstract

Crosstalk between tumor cells and other cells within the tumor microenvironment (TME) plays a crucial role in tumor progression, metastases, and therapy resistance. We present iTALK, a computational approach to characterize and illustrate intercellular communication signals in the multicellular tumor ecosystem using single-cell RNA sequencing data. iTALK can in principle be used to dissect the complexity, diversity, and dynamics of cell-cell communication from a wide range of cellular processes.

---

The TME has emerged as a key modulator of tumor progression, immune evasion, and emergence of the anti-tumor therapy resistance mechanisms^1, 2^. The TME includes a diversity of cell types such as tumor cells, a heterogeneous group of immune cells, and the nonimmune stromal components. Tumor cells orchestrate and interact dynamically with these non-tumor components, and the crosstalk between them is thought to provide key signals that can direct and promote tumor cell growth and migration. Through this intercellular communication, tumor cells can elicit profound phenotypic changes in other TME cells such as tumor-associated fibroblasts, macrophages and T cells, and reprogram the TME, in order to escape from immune surveillance to facilitate survival. Therefore, a better understanding of the cell-cell communication signals may help identify novel modulating therapeutic strategies for better patient advantage. However, this has been hampered by the lack of bioinformatics tools for efficient data analysis and visualization.

Here, we present iTALK (<u>i</u>dentifying and illus<u>t</u>rating <u>al</u>terations in intercellular signaling networ<u>k;</u> https://github.com/Coolgenome/iTALK), an open source R package designed to profile and visualize the ligand-receptor mediated intercellular cross-talk signals from singlecell RNA sequencing data (scRNA-seq) (Fig. 1 and **Online Methods**). We demonstrated that iTALK can be successfully applied to scRNA-seq data to capture highly abundant ligand-receptor gene (or transcript) pairs, identify gains or losses of cellular interactions by comparative analysis, and track the dynamic changes of intercellular communication signals in longitudinal samples. Notably, functional annotation of ligand-receptor genes is automatically added with our curated iTALK ligand-receptor database, and the output can be visualized in different formats with our efficient data visualization tool, which is implemented as part of iTALK. This approach can be applied to data sets ranging from hundreds to hundreds of thousands of cells and is not limited by sequencing platforms. It is also noteworthy that, in addition to studying the TME, iTALK can also be applied to a wide range of biomedical research fields that involve cell-cell communication.

**Figure 1.**
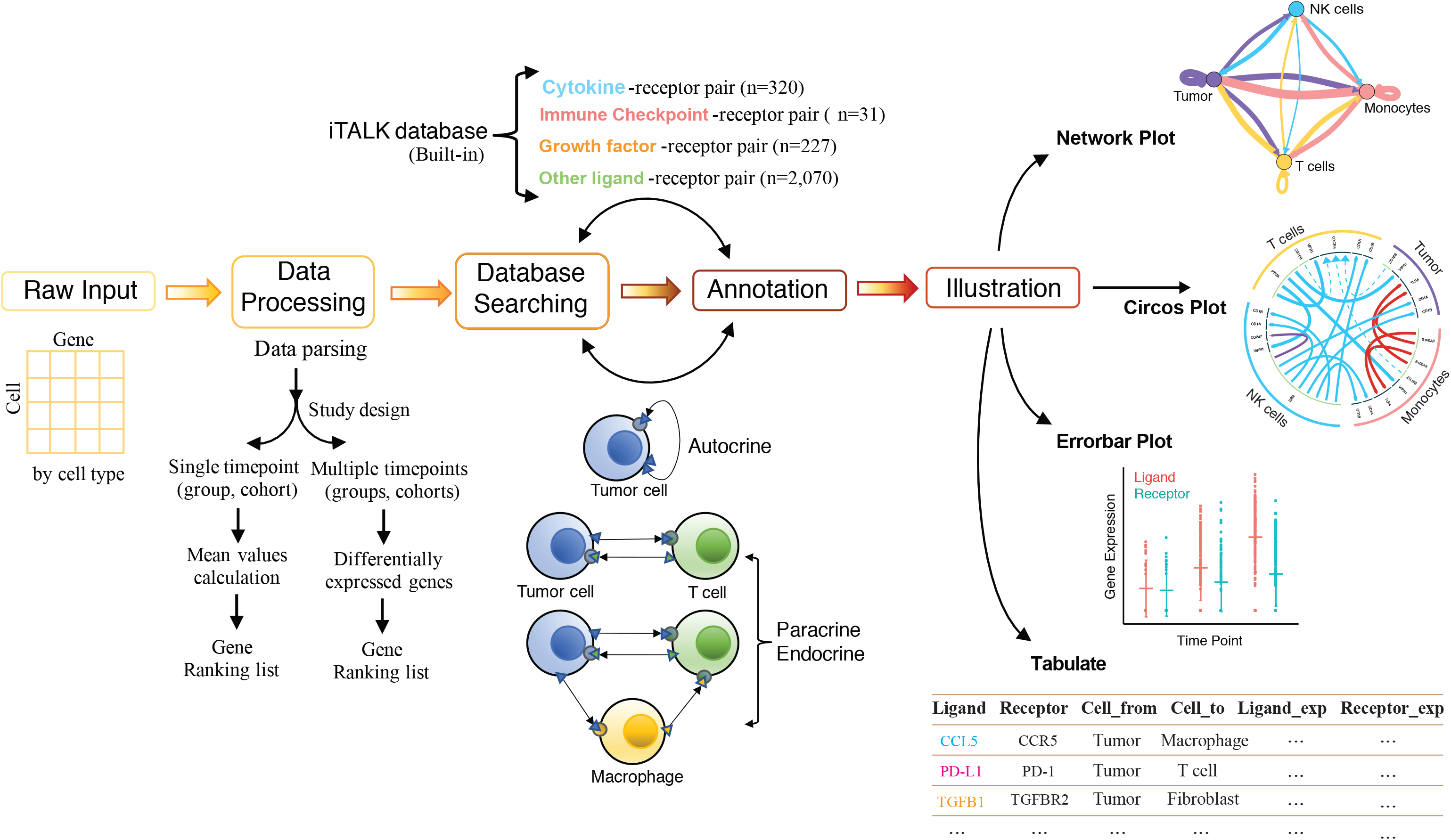
The Bioinformatics workflow of iTALK and illustration formats of iTALK output. iTALK takes cell-gene expression matrix from scRNA-seq as input. The data matrix is processed according to the study design. Known ligand and receptor gene are selected, paired and functional annotated with the built-in ligand-receptor database, and the output can be visualized in different formats with the data visualization tool, which is implemented as part of iTALK.

Ligand-receptor binding is one of the main forms of signal transduction between neighboring and distant cells. To characterize the ligand-receptor mediated intercellular cross-talk, we first manually curated a unique list of ligand-receptor gene pairs based on previous efforts^3–10^ and made this as a built-in database for iTALK. This database collected a total of 2,648 non-redundant and known interacting ligand-receptor pairs, which were further classified into 4 categories based on the primary function of the ligand: cytokines/chemokines, immune checkpoint genes, growth factors, and others (Fig. 1 and **Online Methods**). This database will be updated periodically and user also has the option to input their own gene list or customized gene categories. Given that different cell types may have different receptors for the same ligand and they may induce different response and vice versa, iTALK treats each different ligand-receptor pair as an independent event when analyzing the data. And, because of the functional heterogeneity, the same ligand-receptor binding in different types/phenotypes of cells may trigger distinct cellular response that is independent of their genetic identity. Therefore, scRNA-seq data is an ideal platform to determine which cell type/phenotype expresses which ligand or receptor and to help map out cell-cell communication network more accurately.

iTALK takes the cell-gene expression matrix from scRNA-seq as input. The expression data can be either raw or normalized. The data matrix is parsed and processed subsequently according to the study design. For the dataset that is collected at a single timepoint or for a single group or cohort, iTALK can identify highly expressed ligand-receptor pairs by generating a ranked genes list, followed by searching and pairing ligands and receptors using iTALK database. For the dataset that is collected longitudinally at multiple timepoints or contains multiple genetically/histologically different subgroups or cohorts, iTALK can identify significant changes, i.e. gains or losses of interactions between groups by finding and ranking differentially expressed ligands and/or receptors (Fig. 1). iTALK incorporates commonly used algorithms for batch effects corrections and differential gene expression analysis (**Online Methods**), which also allows the flexibility to select the method that suits the user’s data best.

Notably, iTALK is also an efficient data visualization tool. iTALK output can be visualized in multiple formats including the network plot, the circos plot, the errorbar plot, and the numeric values can be exported as the tabulate format as well (Figs. 1–2). The network plot displays the number of ligand-receptor interactions detected between each two different cell types in the network by labeling the forward (from signaling cell to target cell) and backward signals separately. It also measures the autocrine signals within each individual cell type. The nodes of the network plot are color coded by cell types and the edges are scaled and labelled with the number of interactions between signaling and target cells (Fig. 2a). The circos plot shows the names of each ligand-receptor gene pair and exhibits the direction of each interaction. User has the option to choose which cell type(s), gene category(ies), and the number interactions to be displayed in the circos plot (Figs. 2b-c). The outside ring of circos plot displays cell types, and the inside ring of circos plot shows the details of each interacting ligand-receptor pair. Both are color coded. The lines and arrow heads inside the circos plot are scaled to indicate the relative signal strength of the ligand and receptor, respectively, and different colors and types of lines are used to illustrate various types of possible alterations as shown in Fig. 2. It can demonstrate highly expressed ligand-receptor signals in the scenario of a single timepoint, or group/cohort (Figs. 2b-c), or display the mostly changed (gains or losses) ligand-receptor interactions between two, multiple timepoints, or subgroups/cohorts (Figs. 2d-e). For most ligand-receptor mediated interactions, the ligand appears to have no function unless binding to its receptor. Therefore, we anticipate a loss of interaction (or decrease in interaction) when the expression of a receptor is lost (or decreased), no matter the expression level of its ligand, and vice versa. The errorbar plot is used to demonstrate the dynamic changes of a specific ligand-receptor pair of interest across multiple timepoints (Fig. 2f). All numeric values with annotation information from iTALK can also be exported as a tabulate (Fig. 1), from which, users can sort/filter to select certain interactions for further downstream analysis or customized plotting.

**Figure 2.**
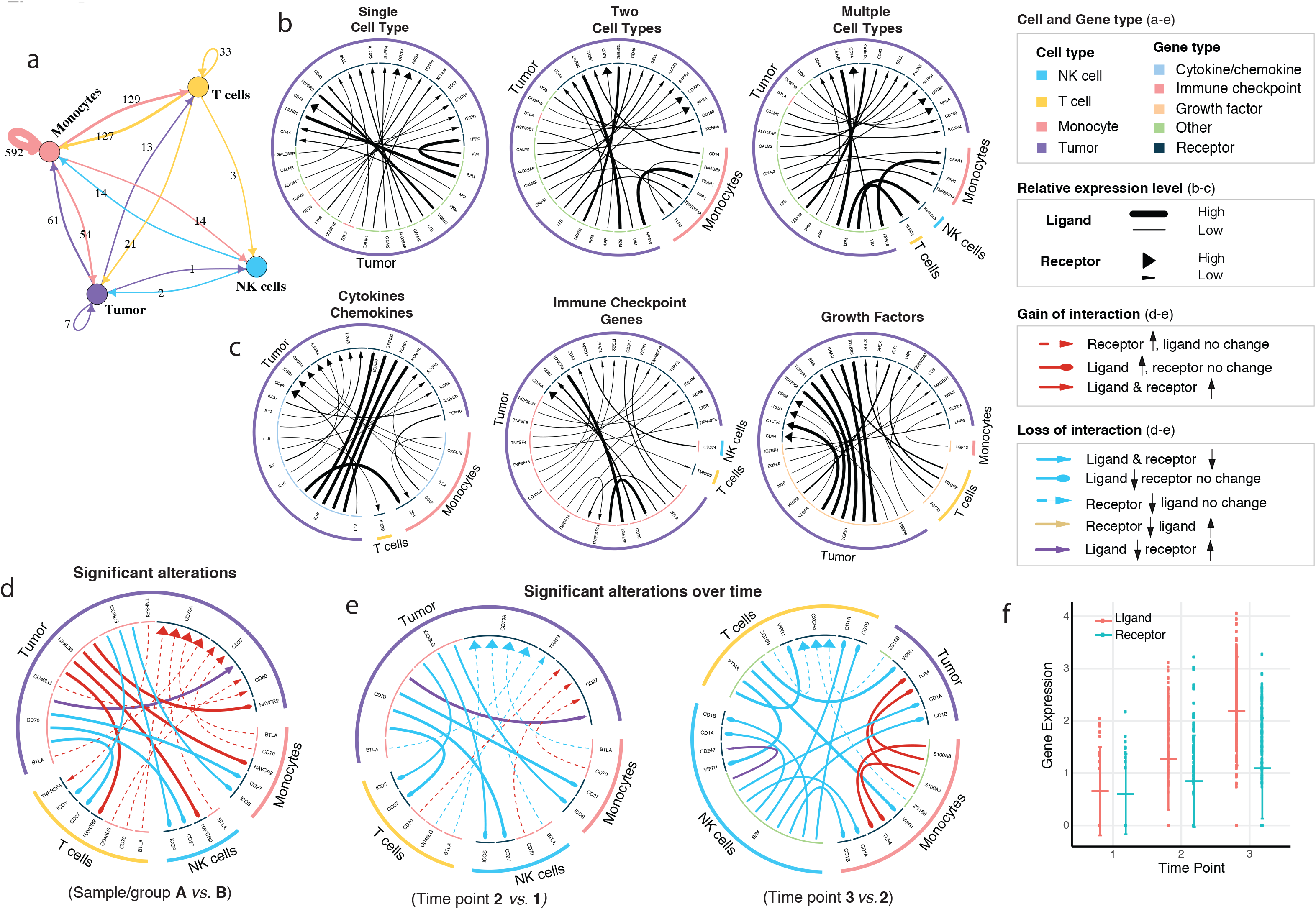
Visualization of iTALK outputs in different formats. **a)** network plot showing the number of ligand-receptor interactions detected between each two different cell types and/or within the same cell type; **b-c)** circos plots for a single timepoint, group or cohort, showing top 20 highly expressed ligand-receptor interactions; **d)** circos plot for a study includes two subgroups or cohorts; or **e)** a longitudinal study, showing significant alterations in cellular interaction. **f)** errorbar plot showing the dynamic changes of a certain ligand-receptor pair across multiple timepoints.

We have successfully applied iTALK to multiple scRNA-seq data sets from both internally generated and publically downloaded. This approach can be applied to data sets ranging from hundreds to hundreds of thousands of cells and is not limited by sequencing platforms. In addition to studying the TME, we anticipate that iTALK can also be applied to a wide range of biomedical research fields that involve cell-cell communication to help dissect the complex intercellular signaling networks.

## METHODS

Method, including statements of data availability and any associated accession codes and references, are available in the online version of the paper.

## ACKNOWLEDGEMENTS

This work was supported by the start-up research funds kindly awarded to L.W. by the Department of Genomic Medicine, Division of Cancer Medicine of MD Anderson Cancer Center. This work was also supported in part by the U.S. Department of Defense (DOD) grants-CA160445 to J.A., and the generous philanthropic support to the MD Anderson B Cell Lymphoma Moon Shot Project (M.W.).

## AUTHOR CONTRIBUTIONS

Conception and design: L.W. Development of methodology: L.W., Y.W., R.W. Sample collection and processing: S.S., J.A., C.J., M.W. Analysis and interpretation of data: Y.W., L.W., S.Z., G.H., A.F. Writing, review and/or revision of the manuscript: L.W., Y.W., J.A., M.W., A.F. Study supervision: L.W.

## ONLINE METHODS

### Database and annotation

The built-in database is manually curated from previous efforts^3–10^. The gene lists were combined and duplicates were removed to make a non-redundant ligand-receptor list. This database contains a total of 2,648 unique ligand-receptor interacting pairs. We further classified them into four categories: cytokine/chemokine (n=320), immune checkpoint (n=31), growth factor (n=227) and others (n=2,070), based on the known function of the ligand. We anticipate to update the database periodically to keep it up-to-date. In addition to the built-in database, user also has the option to input their own ligand-receptor gene list. They can also customize the gene categories as needed.

### Input data format, data parsing, filtering, batch effects removal

The input data should be a cell-gene expression matrix with cell type or phenotype annotated for each cell. Raw data should be filtered based on QC metrics. The default criteria are to filter out genes detected in <3 cells and cells where <200 genes had nonzero counts. Low-quality cells where >15% of the counts derived from mitochondrial genome will also be discarded. For sequencing data with possible batch effects, scde^11^ and monocle^12^ (incorporated into iTALK) can be applied to remove them.

### Identifying significant interactions

We integrated multiple algorithms for identifying significant interactions (differentially expressed ligands and receptors) including commonly used R packages: DESeq2^14^, scde^11^, monocle^12^, DEsingle^15^, edgeR^16^ and MAST^17^. User has the flexibility to select the method that suits their data set best. All these methods require the input data to be raw, without any filtering or normalization. Considering single cell sequencing data sometimes can have thousands or tens of thousands of cells, we also provided Wilcoxon method to speed up data processing. The input data for Wilcoxon method can be both raw or processed. After finding the highly expressed or differential expressed genes, these genes will be matched and paired using our ligand-receptor database to find significant ligand-receptor interactions. To identify alterations in singling interactions, the expression levels of both ligand and receptors are evaluated.

We define the gain of interaction if either a ligand (or receptor) gene upregulated and its paired gene upregulated or remains no change. We define the loss of interaction if either the ligand (or receptor) gene downregulated, no matter the expression level of its paired gene.

### Illustration

The network plot, circos plot and errorbar plot are all color coded and user has the option to choose their favorite colors (e.g. for cell types, gene categories, strength and direction of changes, etc.). In addition, use has the options to choose the number of cell types (tumor cells, T cells, NK cell, macrophages, fibroblasts, etc.) or gene categories (cytokine/chemokine, immune checkpoint, growth factor, and others), and the number of interactions (top 30, 50, 100, or all) to be displayed in the circos and network plots. The network plot is generated based on the R package igraph^18^; circos plot is based on package circlize^19^, and errorbar plot is based on ggplot2^20^.

### Example datasets

Our example datasets were generated internally. The data were filtered and normalized by Seurat^13^. The cell types are defined with known markers. Briefly, B cells were identified using CD79A, CD79B and MS4A1; T cells were identified by CD3D/E; CD8 T cells were identified using CD3D/E and CD8A; NK cells were classified by CD45, NKG7, GZMA; and monocytes were identified using CD68, CD163, S100A8/9. Wilcoxon method was applied to find significant interactions between different groups.

## Code availability

The code and built-in database are publicly accessible on GitHub at https://github.com/Coolgenome/iTALK

